# Microtubules deform the nucleus and force chromatin reorganization during early differentiation of human hematopoietic stem cells

**DOI:** 10.1101/763326

**Authors:** S. Biedzinski, L. Faivre, B. Vianay, M. Delord, L. Blanchoin, J. Larghero, M. Théry, S. Brunet

**Author notes:** Correspondence should be sent to and.

## Abstract

Hematopoietic stem cells (HSC) can differentiate into all hematopoietic lineages to support hematopoiesis. Cells from the myeloid and lymphoid lineages fulfill distinct functions with specific shapes and intra-cellular architectures. The role of cytokines in the regulation of HSC differentiation has been intensively studied but our understanding of the potential contribution of inner cell architecture is relatively poor. Here we show that large invaginations are generated by microtubule constraints on the swelling nucleus of human HSCs during early commitment toward the myeloid lineage. These invaginations are associated with chromatin reorganization, local loss of H3K9 trimethylation and changes in expression of specific hematopoietic genes. This establishes the role of microtubules in defining the unique lobulated nuclear shape observed in myeloid progenitor cells and suggests that this shape is important to establish the gene expression profile specific to this hematopoietic lineage. It opens new perspectives on the implications of microtubule-generated forces, in the early specification of the myeloid lineage.

## INTRODUCTION

Hematopoietic stem cells (HSCs) sustain hematopoietic lineages homeostasis throughout life (Orkin and Zon, 2008). A cell’s engagement toward specific differentiation pathways is the outcome of the interplay of numerous cytokines (Notta et al., 2016; Velten et al., 2017) as well as physical cues associated with its interactions with the extra-cellular matrix and with stromal cells in the bone marrow (Crane et al., 2017; Pinho and Frenette, 2019). Whether and how the transduction of these signals is mediated by specific intra-cellular reorganization of cytoskeleton networks and organelle morphologies is not yet known; although these parameters have well established roles in the regulation of other lineages (Uhler and Shivashankar, 2017).

For example, the fate of mesenchymal stem cells strongly depends on the architecture of the acto-myosin network (Kilian et al., 2010; McBeath et al., 2004). Substrate stiffness and adhesiveness promotes the formation of large focal adhesions and cytoplasmic acto-myosin bundles, the contraction of which deforms the nucleus (Buxboim et al., 2014; Khatau et al., 2009), induces chromatin remodeling (Makhija et al., 2015; Versaevel et al., 2012) and the shuttling of transcription factors (Dupont et al., 2011; Miralles et al., 2003) that further impact gene expression and cell identity (Alam et al., 2016; Gupta et al., 2012; Närvä et al., 2017; Tajik et al., 2016). However, similar mechanisms are unlikely to impact the fate of HSCs because these cells are poorly adhesive and thus are unable to assemble focal adhesion and contractile bundles. The contraction of the cortical actin network can modulate cell shape and the symmetry of HSC division (Shin et al., 2013, 2014) but has not been shown to deform cell nucleus as in the case of mesenchymal stem cells. However, differentiated blood cells can display highly deformed nuclei, notably neutrophils (Bainton et al., 1971; Carvalho et al., 2015; Hoffmann et al., 2007). It is not known whether such deformations are instrumental in the regulation of the differentiation process, when they are initiated during cell differentiation, or what is mechanism that drives them.

To tackle these questions and better understand the early structural rearrangements of HSC, we investigated the architecture of primary human HSCs as they commit either to the lymphoid or the myeloid lineages.

## RESULTS

We investigated the nucleus shape at various early stages of human HSC differentiation, namely stem cells, myeloid progenitors and lymphoid progenitors. We used human neonatal cord blood-isolated and FACS-purified cell populations. The surface-marker signature of stem cells was considered to be CD34+/CD38-, whereas differentiated progenitors were CD34+/CD38+. Progenitors engaged in myeloid and lymphoid pathways were considered to be, respectively, CD33+ and CD19+ (Notta et al., 2016)(Figure 1A). Simple morphometric analysis revealed that nuclei displayed distinct and specific sizes and shapes at each stage of early hematopoietic differentiation. To measure this, sorted cells were fixed, immunostained and imaged with highly resolved confocal microscopy. A quantitative analysis of nucleus volume and surface curvature was performed using 3D reconstructions (see Material and Methods) (Figure 1B and C). In particular, the ratio of concave surface areas on the total surface was calculated as a proxy for the degree of invagination of the nucleus. Stem cells displayed rather small and round nuclei, characterized by two populations with volumes around 120 and 190 µm^3^ and a low mean invagination ratio of 8% for both. The nuclei of lymphoid progenitors were also small (120 µm^3^ on average) and spherical with a mean invagination ratio similar to those of HSCs (Figure 1B and C). By contrast, the nuclei of myeloid progenitors were approximately twice the size (250 µm^3^ on average). In addition, they displayed large deformed areas and invaginations. Accordingly, their invagination ratio of 15% was almost 2-fold greater than for stem cells or lymphoid progenitors (Figure 1B and C). This suggested that nucleus deformation is involved in the regulation of the myeloid differentiation process.

**Figure 1.**
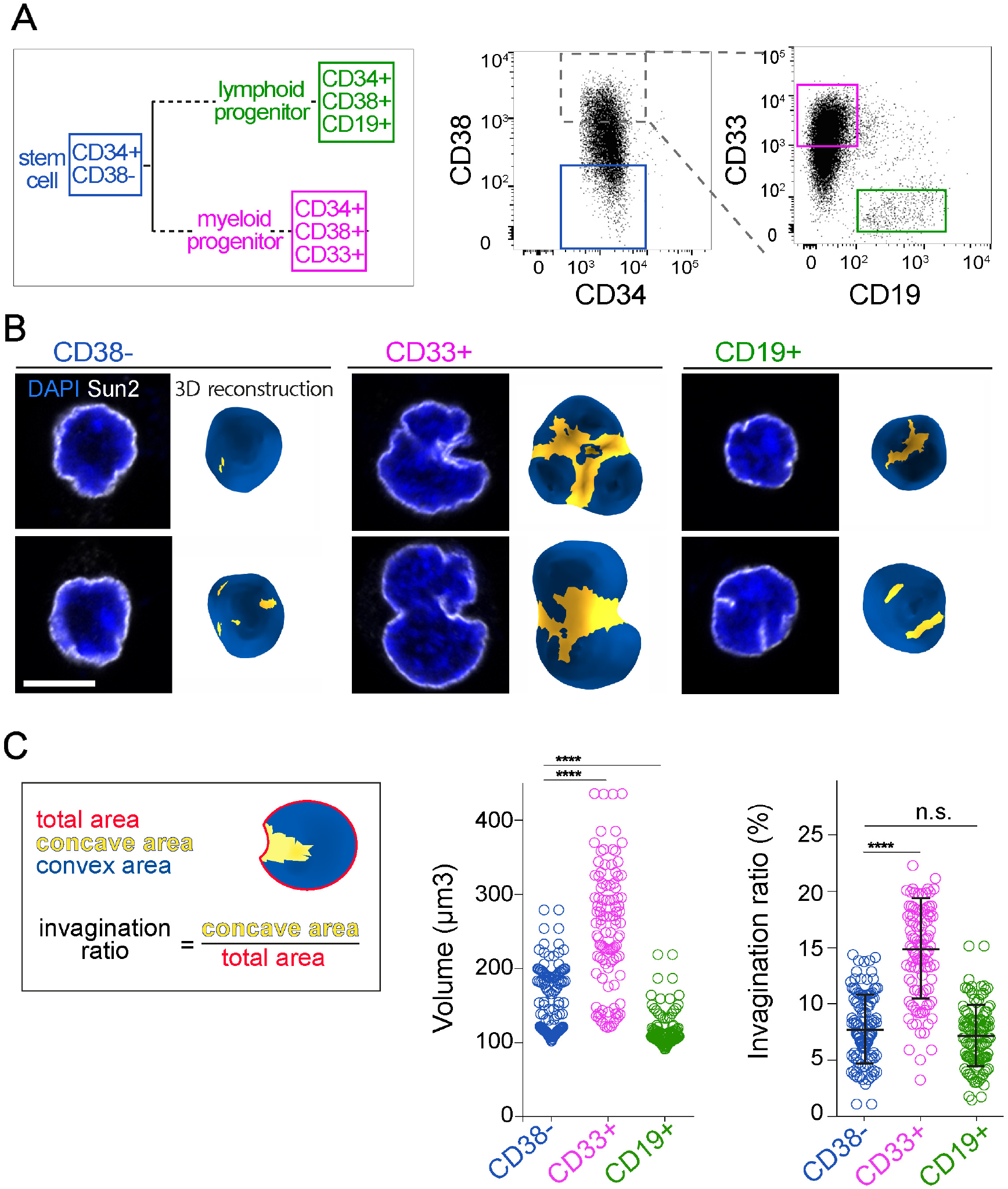
Nucleus morphologies of stem cells and progenitors. A) Left panel: experimental strategy. Hematopoietic stem cells (HSC) and progenitors were characterized and isolated using specific sets of surface markers. All cells express CD34. Upon differentiation, progenitors start to express CD38 in contrast to HSCs (blue). Progenitors engaged in the myeloid (magenta) and lymphoid (green) pathways express respectively CD33 and CD19 markers. Right panel: FACS gating strategy to isolate HSCs (further referred as CD38-(blue box), and cells engaged in myeloid and lymphoid differentiation pathways (referred to as CD33+ [magenta box] and CD19+ [green box], respectively). B) For each population, equatorial Z stacks of two representative nuclei are shown in the left panel (chromatin is shown in blue (DAPI staining) and nuclear envelop in white (Sun2 staining). Scale bar: 5 µm. The corresponding 3D reconstructions are presented in the right panel. Convex and non-convex surfaces are shown in blue and yellow, respectively. C) Nucleus-invagination ratio is defined as the ratio of the non-convex area over the total nucleus surface area. Nuclei are larger and deformed in CD33+ (magenta, n=106) than CD38- and CD19+ cells (blue [n=120] and green [n=115], respectively; 3 independent experiments; ***: p<0.001. ****: p<0.0001, Mann-Whitney test; n.s: non significant).

We then investigated differences in the intra-nuclear and cytoplasmic cytoskeleton networks between early myeloid progenitors and stem cells that could be responsible for the observed deformations. The content of laminA/C over laminB was higher in myeloid progenitors than in HSCs (Supplemental Figure S1A and B) arguing against a global softening of the nuclear membrane as the cells started to differentiate (Makhija et al., 2015; Swift et al., 2013). In both cell types, actin was mainly present at the cell cortex (Figure 2A) but was not found on the nucleus surface. By contrast, microtubules were closely associated with the nucleus (Figure 2A and B). In myeloid progenitors, numerous microtubules appeared to form bundles within nuclear invaginations (Figure 2A and B). Microtubules were found to converge at one of the larger invaginations, where the centrosome (the main microtubule organizing center) was located (Figure 2B). This could be quantified by measuring the relative position of the centrosome to the nucleus center and the cell periphery (Figure 2C). In contrast to HSCs where the centrosome was located close to cell periphery, in myeloid progenitors, the centrosome was buried deep in nuclear invagination, suggesting an active role of the centrosome-microtubule network in the deformation of the nucleus.

**Figure 2.**
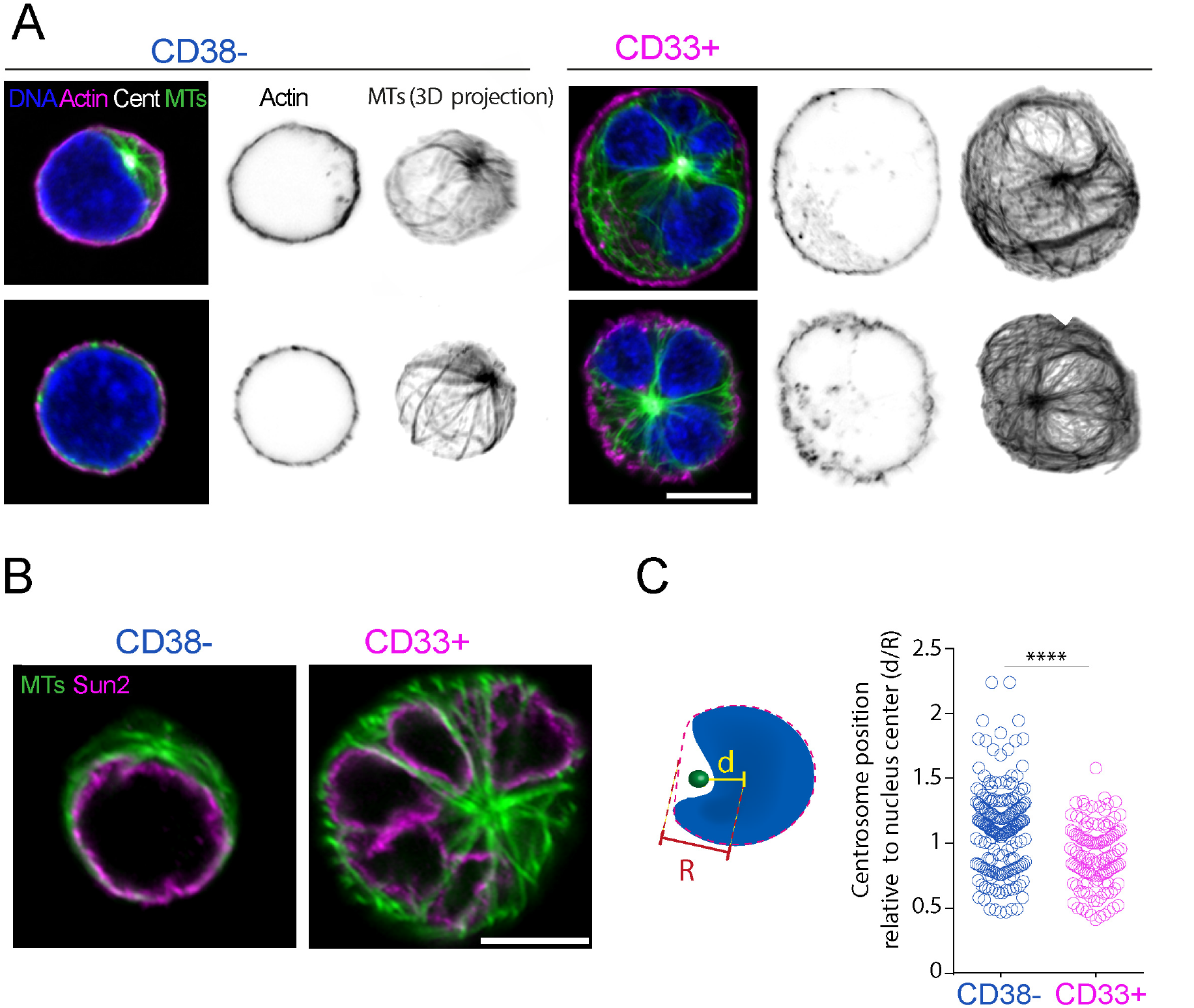
Cytoskeletal organization of stem cells and myeloid progenitors. A) For each population (CD38- and CD33+ cells), Z stacks of two representative cells are shown in the left panel. Microtubules are shown in green, actin in magenta, centrosome in white and chromatin in blue. In the middle panel, inverted image the corresponding actin staining is shown. In the left panel, the corresponding 3D projection of microtubules is presented as inverted image. Scale bar 5 µm. B) Microtubules are tightly associated with the nucleus. Z stacks of representative CD38- and CD33+ cells are shown. Microtubules are shown in green, nucleus envelope (Sun2 immuno-staining) in magenta and chromatin in blue. Scale bar 5 µm. C) As schematized, centrosome-to-nucleus center distance (d) over nucleus center-to-nucleus convex envelope (R) ratio is calculated to extract relative position of the centrosome to the center. This parameter is significantly lower in myeloid progenitors (n=129) than in HSCs (n=200; 3 independent experiments; ****; p<0.0001. Mann-Whitney test) indicating that centrosome becomes internalized within an invaginated region upon differentiation.

Microtubule disassembly seemed like a straightforward way to challenge the role of microtubules in nuclear deformation. However, a 3 hour-long incubation with nocodazole, which fully depolymerized microtubules, had no impact on the nucleus deformations (Supplemental Figure S1C). This indicated that nuclei of myeloid progenitors had undergone plastic deformations that were introduced earlier in the course of differentiation. Therefore, to further test the role of microtubules, it was necessary to act upstream of the invagination process to prevent it. We first determined the exact time window during which nucleus was reshaped as cells progressed through differentiation. To achieve this, isolated HSCs were grown in culture medium supplemented with SCF, G-CSF and IL3 to promote their differentiation into myeloid progenitors (Donaldson et al., 2001; Faivre et al., 2016) and fixed at various time points. (Figure 3A). After 48 h of culture, nuclei had enlarged and displayed significant deformation, exhibiting kidney-like shapes with shallow invaginations tightly associated with centrosomes and microtubules (Figure 3B). After 72 h of culture, nuclei were substantially deformed with deep invaginations containing microtubule bundles (Figure 3B). The nucleus volume and the invagination ratio at that stage were similar to those measured in myeloid progenitors freshly isolated from cord blood (Figure 3C). Therefore the differentiation of HSCs for 3 days in culture was sufficient to recapitulate the nucleus expansion and deformation characteristics of primary myeloid progenitors.

**Figure 3.**
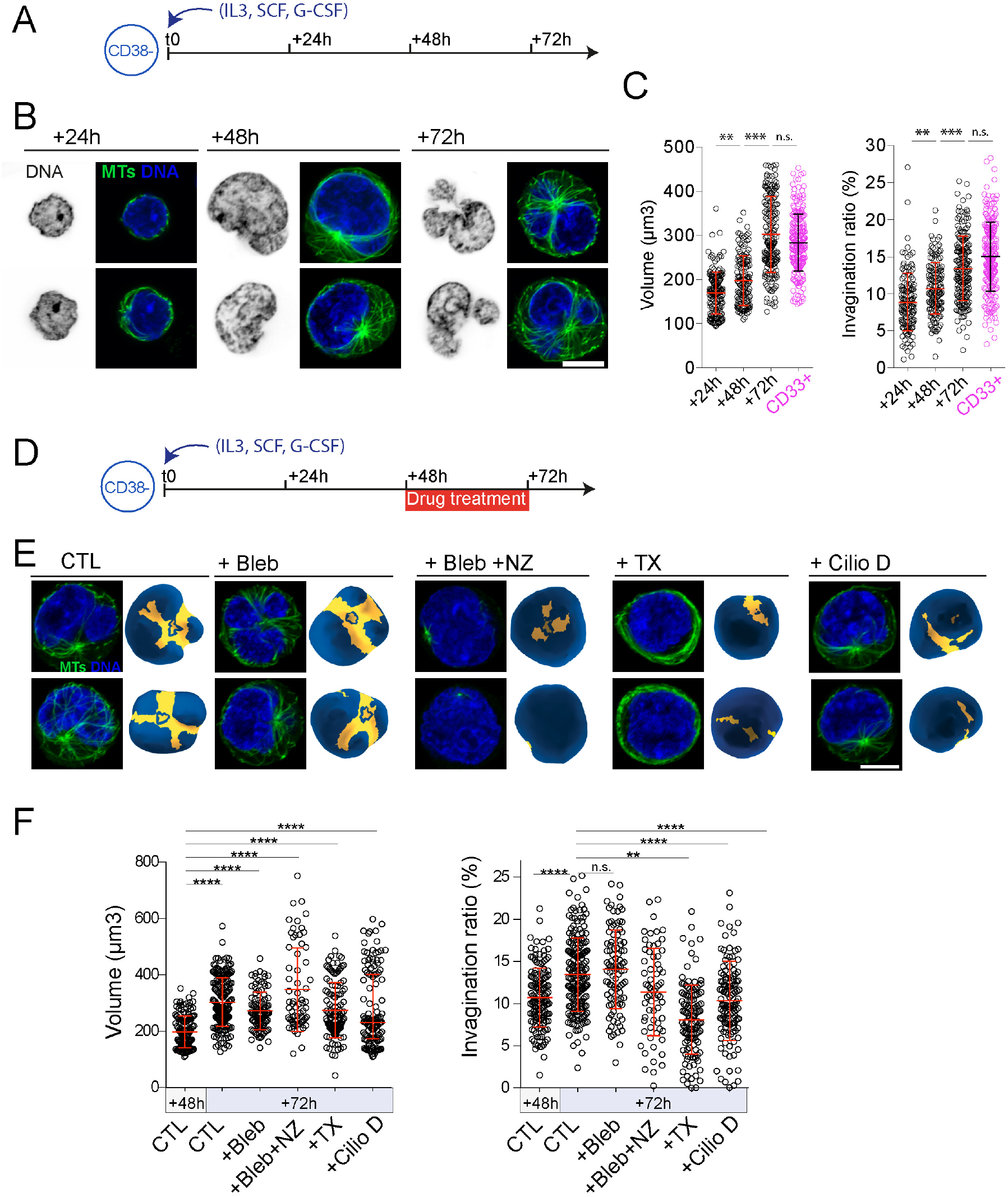
Microtubules reshape the nucleus during HSC differentiation *in vitro*. A) Experimental set-up: isolated CD38-cells are cultured and differentiated upon addition of IL3, SCF and G-CSF cytokines. Cells are collected and analyzed at 24, 48 and 72 hours after cytokine addition. B) The nucleus is progressively deformed over time in culture. For each time point, equatorial Z stacks of two representative cells are presented. Left panel: inverted image of chromatin (DAPI). Right panel: microtubules are shown in green and chromatin in blue. Scale bar: 5 µm. C) Nucleus volume (left panel) and invagination ratio (right panel) increase to attain levels similar to those of freshly isolated CD33+ cells (24 h, n=165, 4 donors; 48 h, n=143, 3 donors; and 72h, n=192, 4donors; CD33+, n=227, 5 donors; n.s: non significant; ***: p<0.001, ****: p<0.0001; Mann Whitney test). D) Experimental set-up: indicated drugs were added to culture medium between 48 and 72 hours. Cells were fixed and analyzed after 72 hours of culture. E) For each condition (CTL, untreated cells; +Bleb, blebbistatin-treated cells; +Bleb+NZ, blebbistatin and nocodazole-treated cells; +TX, taxol-treated cells; and +CilioD, ciliobrevin-D–treated cells), Z stacks of 2 representative cells are presented in the left panel. Microtubules are shown in green, chromatin in blue. In the right panel, the corresponding 3D reconstructions of DAPI staining are shown. Convex and non-convex surfaces are shown in blue and yellow, respectively. Scale bar 5 µm. F) Quantifications of nuclear volume (left panel), invagination ratio (right panel) in the indicated conditions. Non-treated cells at 48 hours (in blue, n=137, 3 donors) and 72 hours (n=143, 3 donors); and blebbistatin-treated cells (n= 96/3 donors), blebbistatin and nocodazole treated cells (n=66, 2 donors), taxol-treated cells (n=122, 3 donors), and ciliobrevin-D–treated cells (n=128, 3 donors).

Given the most substantial nucleus deformations of HSCs occurred between Day 2 and Day 3 in culture, this was selected as the appropriate time window to interfere with microtubule network organization (Figure 3D). As in many other cell models, microtubule depolymerization induced the contraction of the actin cortex (Kolodney and Elson, 1995; Paluch et al., 2005). This side effect, which could indirectly impact nucleus shape (Supplemental Figure S1C), was compensated by the addition of blebbistatin to nocodazole (Figure 3E). Hence, in the absence of microtubules, the nucleus of differentiating HSCs expanded but did not deform (Figure 3F). A control experiment showed that blebbistatin alone was not sufficient to prevent nucleus deformation (Figure 3E and F). The active role of microtubules was further investigated by stabilizing them with taxol or by inhibiting dynein, a motor mediating nucleus-microtubule interaction (Firestone et al., 2012; Salina et al., 2002)(Figure 3E). Taxol-treated microtubules formed a dense array along the cell cortex, which was no longer in contact with the nucleus (Figure 3E). In both conditions, nuclear expansion still occurred but nuclear deformations were completely impaired (Figure 3F). Taken together, these experiments demonstrated that microtubules and dyneins were responsible for nucleus invaginations during HSC differentiation into myeloid progenitors.

To investigate whether nuclear deformation could modulate gene expression and thereby HSC differentiation, we analyzed its relationship with the spatial organization of chromatin, and more specifically with the localization of dense heterochromatin, which acts as a repressor of gene expression (Mattout et al., 2015). We first assessed the spatial distribution of histone H3 lysine 9 trimethylation (H3K9me3), as a marker of constitutive heterochromatin (Becker et al., 2016)(Bannister et al., 2001). In freshly isolated HSCs, and in lymphoid progenitors, H3K9me3 was homogeneously distributed along the entire peripheral margin of the nucleus (Figure 4A), as previously described (Djeghloul et al., 2016; Ugarte et al., 2015). By contrast, in freshly isolated myeloid progenitors, H3K9me3 distribution was more heterogeneous: foci were detected in the nucleoplasm, and at the peripheral margin of the nucleus, the density of H3K9me3 foci was lower in the invaginated regions than the other (convex) regions (Figure 4A). Moreover, the difference of H3K9me3 distribution between the HSC state and the myeloid-progenitor state was also observed in the differentiation of HSCs into myeloid progenitors in culture (Figure 4B). The quantification of H3K9me3 distribution by linescan density measurements at the peripheral margin of the nucleus of cultured HSCs (Figure 4C and D and Supplemental Figure S2A), revealed a correlation between H3K9me3 density and curvature (Figure 4D). Hence, H3K9me3 density was lower with negative (inward) curvature, and this association was more acute after 3 days of HSC differentiation toward myeloid lineage (Figure 4E and Supplemental Figure S2B). This relationship was further supported by quantifying the dispersion of H3K9me3 signal intensity at the peripheral margin of the nucleus in cultured HSCs (Fig 4F and Supplemental Figure S2C). The signal was generally homogeneous after 24 h in culture, but was more heterogeneous after 48h, and attained a degree of dispersion after 72h that was similar to that observed in freshly isolated myeloid progenitors (Figure 4F). In contrast to myeloid progenitors, a heterogeneous dispersion of H3K9me3 was not observed in lymphoid progenitors (Supplemental Figure S2D). These trends accurately followed the extent of nucleus deformation as the cells differentiated (Figure 3C).

**Figure 4.**
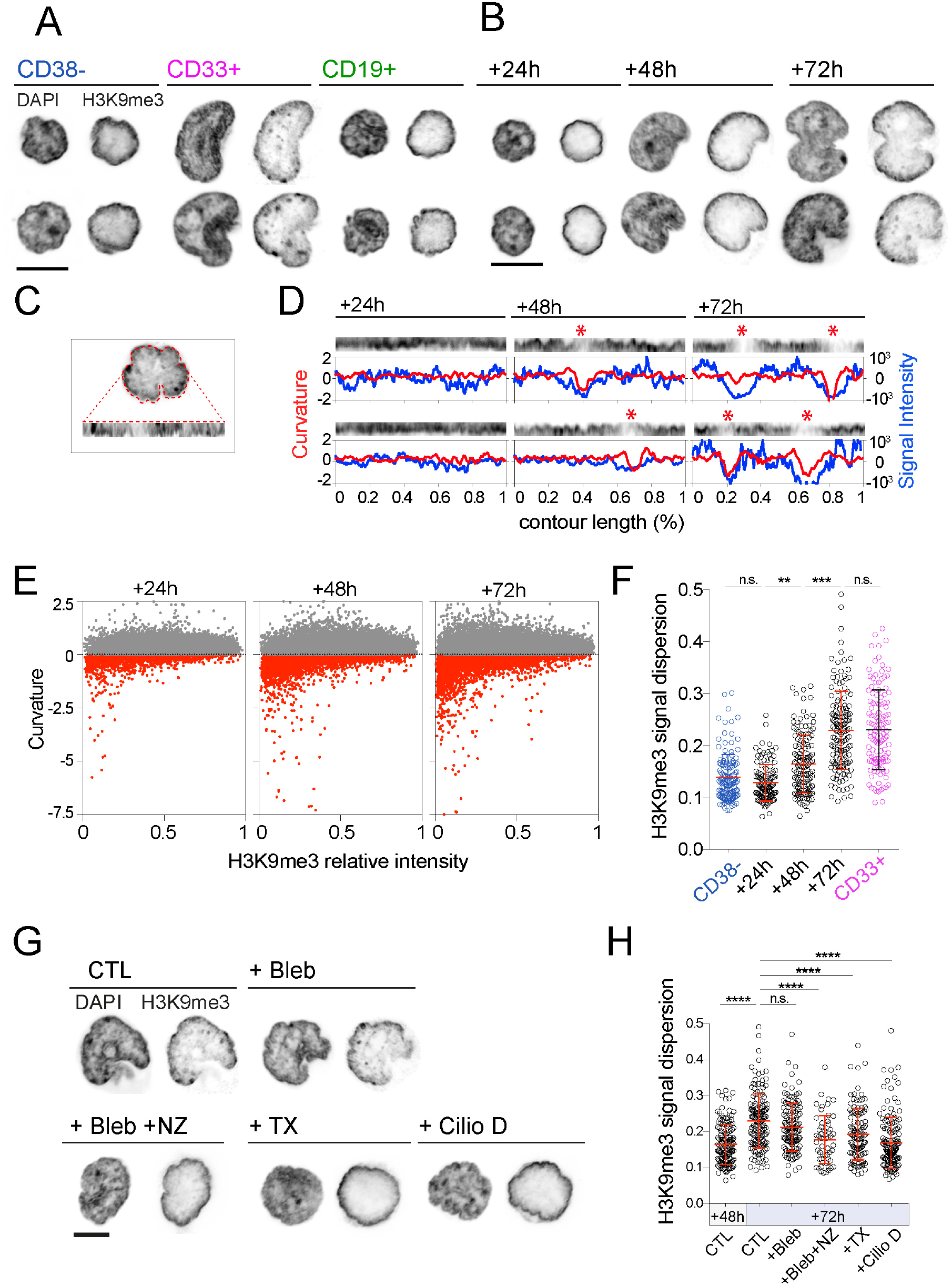
HSC differentiation is associated with microtubule-dependent heterochromatin redistribution from the nucleus periphery. A) Spatial distribution of H3K9me3 in CD38-, CD33+ and CD19+ cells. For each population, two representative nuclei are presented. Inverted images of equatorial Z plane of DAPI (left panel) and H3K9me3 (right panel) are shown. Scale bar: 5 µm. B) Spatial distribution of H3K9me3 at the indicated time points of differentiation of CD38-cells. For each population, two representative nuclei are presented. Inverted images of equatorial Z plane of DAPI (left panel) and H3K9me3 (right panel) are shown. Scale bar: 5 µm. C) For each cell, the raw DAPI image at the equatorial Z stack is used to extract the nucleus contour and measure H3K9Me3 intensity (in arbitrary units [a.u]) along the contour (lower row). D) H3K9me3 loss from the nucleus periphery occurs locally in nuclear invaginations. Line scans of H3K9me3 signal along nucleus contour of 2 representative cells are presented in the upper panel for each time point. Asterisks indicate the local loss of H3K9me3. The corresponding variations in H3K9Me3 intensity and curvature are plotted in blue in red, respectively. Negative curvature values correspond to local nuclear invaginations. E) Local curvature of the nucleus envelope as a function of H3K9Me3 local intensity at the indicated time points of differentiation. Positive and negative curvatures are shown in grey and red, respectively. The absence of curvature is highlighted as a dashed line. During differentiation, negatively curved domains increase in number and size and display low H3K9me3 intensities. F) H3K9me3 redistribution during differentiation. H3K9me3 signal dispersion at the nucleus periphery is low in CD38-(n=137, 3 donors) indicating that H3K9me3 is homogenously distributed at the nucleus periphery. Upon differentiation, H3K9me3 signal dispersion remains homogenous after 24 hous in culture (n=115, 3 donors), but progressively becomes heterogeneous (48 h: n=139, 3 donors) to attain a level at 72 h (n= 148, 3 donors) similar to that in freshly collected CD33+ cells (n=124, 3 donors), indicative of H3K9me3 redistribution from the nucleus periphery. G) Microtubule perturbations impair H3K9me3 redistribution during HSC differentiation. Microtubule-modifying agents were added to culture medium between 48 and 72 hours. Cells were fixed and analyzed after 72 hours of culture. For each indicated treatment, one representative nucleus is presented. Inverted images of equatorial Z stacks of DAPI (left panel) and H3K9me3 (right panel) are shown. Scale bar: 5 µm. H) H3K9me3 signal dispersion significantly increases in non-treated cells between 48h (n=139/3 donors) and 72h (n=148/3 donors) of culture. At 72h, the dispersion is similar in non-treated and blebbistatin-treated cells (n=124/ 3 donors). By contrast, H3K9me3 signal dispersion is significantly reduced compared with non-treated cells after treatment with nocodazole and blebbistatin (n=55, 2donors), taxol (n= 109, 3 donors) and ciliobrevin-D (n=144, 3donors) to a level attained by non-treated cells at 48 hours, demontrating that microtubules affect H3K9me3 redistribution during HSC differentiation.

Chromatin reorganisation can participate in neutrophil nucleus reshaping (Zhu et al., 2017). Hence the local loss of heterochromatin could initiate nuclear invagination. Alternatively, it could result from it. To distinguish these two possibilities we prevented nuclear invagination, by interfering with microtubule-nucleus interaction, and then monitoring H3K9me3 distribution along the nuclear membrane. The three treatments described above, blebbistatin/nocodazole, taxol or ciliobrevin D, were applied during the critical day-2 to day-3 period, during which nuclei in differentiating HSCs undergo the invagination process. At 72 hours of culture, non-treated cells and control cells treated with blebbistatin exhibited the expected heterogeneous H3K9me3 patterns associated with high signal dispersion values (Figure 4G and H). By contrast, cells treated with drugs against microtubules, displayed minor invaginations of the nuclei (Figure 3F and 4G) and no significant increase of H3K9me3 signal dispersion (Figure 4H). These results showed that microtubule-induced invaginations were required for the loss of heterochromatin in these regions.

We finally tested whether the heterochromatin remodeling that resulted from nuclear invagination could actually impact the process of myeloid differentiation. The investigation of the link between microtubule-controlled nucleus deformation and heterochromatin redistribution and myeloid differentiation was complicated by the fact long-term interference of microtubules could impair cell cycle progression. Therefore in cultured HSCs (from three donors), microtubules were perturbed between day 2 and day 3 by the addition of the dynein inhibitor ciliobrevin D, and the effect on the transcriptome was assessed (Figure 5 and Supplemental Table 1). In total, 123 genes were found to be significantly up- or down-regulated with ciliobrevin D treatment (absolute log2 fold change higher than 1.8 and a false discovery rate lower than 0.05). Strikingly, among the 38 genes that were significantly down-regulated, eight had been previously defined as gene signatures of cells of myeloid lineages, notably erythroblasts, including CD14+ and CD66b+ lineages (*BSG; CPVL; TLY96; QPCT; ZER1; SLC2A11: IL6R; IFIT3*) (Watkins et al., 2009). Among the up-regulated genes, two genes were erythoblast-specific (*EPB42, FKBP4*), one gene was CD66b+ lineage-specific (*TM6SF*) but none were associated with the CD14+ lineage. Importantly, six of the up-regulated genes were associated with the megakaryocyte differentiation pathway (*MED12L, CTDSPL, FUT8, ABCC4 FCER1A and CPA3*), which separates from the myeloid pathway in early stages of hematopoiesis (Notta et al., 2016). These results show that, although microtubule perturbation was restricted to a small period within the HSC differentiation process, it was sufficient to impair HSC differentiation into myeloid progenitors.

**Figure 5.**
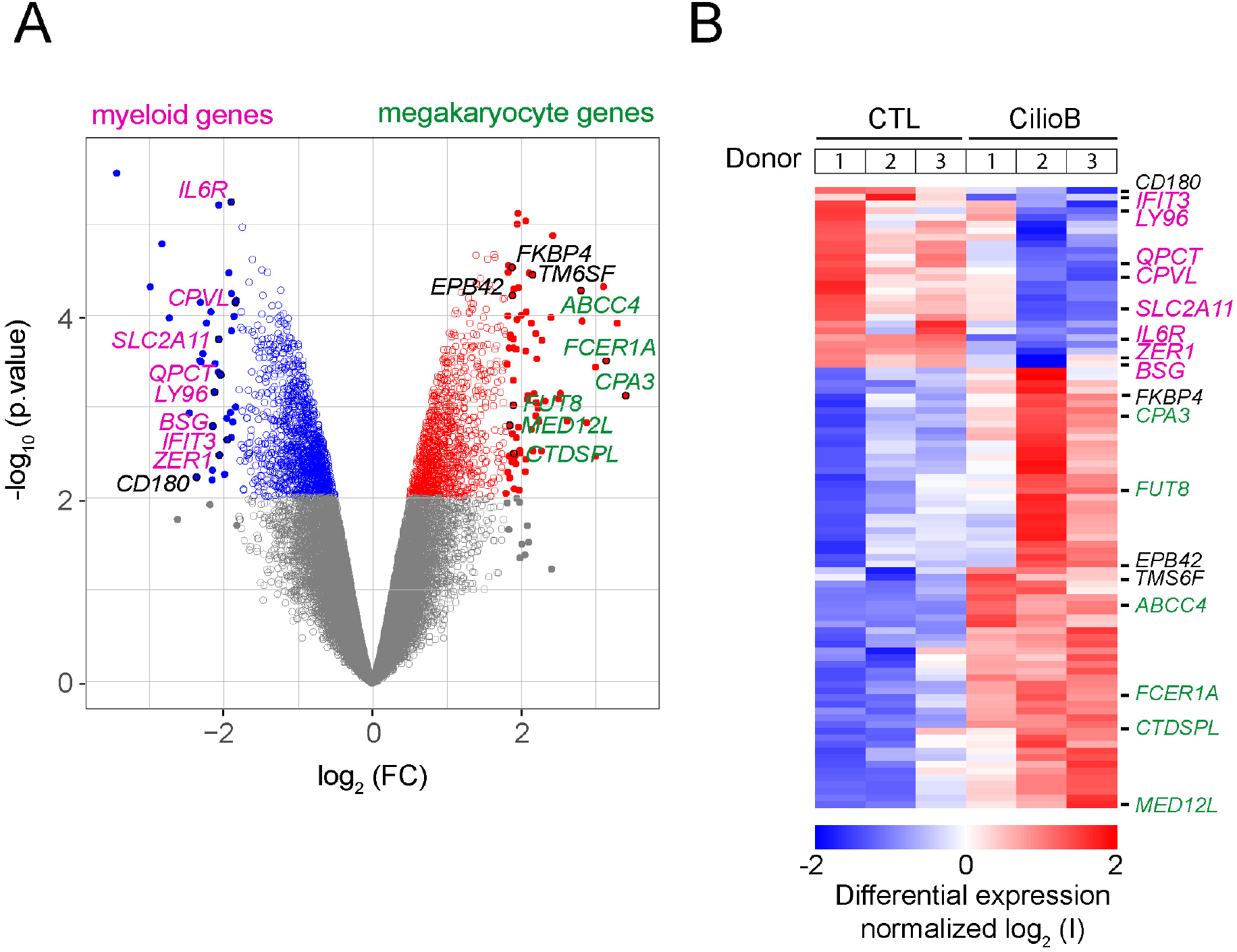
Microtubules impact the trancriptome of myeloid differentiating cells. A) Microtubules perturbations impact the trancriptome of differentiating cells. Volcano-plot representation of down-regulated (blue) and up-regulated genes (red) in non-treated/control cells relative to ciliobrevin-D treated cells. Each circle represents the fold change (FC) of the quantity of mRNA and the significance of the change based on three independent replicates. Expressed genes with a false discovery rate (FDR) lower than 0.05 are shown in grey, whereas genes with a FDR greater than 0.05 are shown in blue or red. Genes significantly mis-regulated i.e. with an absolute value of FC higher than 1.8 are shown as full disks. Names of genes of interest are indicated within the graph. Genes that are in the signature of the myeloid-differentiation pathway are shown in magenta. Genes that are the signature of the megakaryocyte-differenciation pathway are shown in green. B) Heatmap representation and hierarchical clustering of gene expression changes between non-treated and ciliobrevin-D–treated cells collected from 3 donors in each condition. The color code corresponds to a log_2_ scale of the differential expression levels.

In conclusion, our study shows that in contrast to the lymphoid differentiation pathway, HSC commitment to the myeloid pathway involves the formation a large nuclear membrane invaginations and local reorganization of heterochromatin. During this process microtubules act as a mechano-regulator of stem cell identity. They form bundles that encircle the nucleus and resist chromatin expansion. This generates nuclear invaginations in which the density of constitutive heterochromatin is reduced. These structural rearrangements can affect the expression of genes, some of which are involved in progenitor identity and function.

## DISCUSSION

The pioneering observations of Paul Ehrlich revealed the highly lobulated shapes of some mature blood cells (Ehrlich and Lazarus, 1900). In the case of neutrophils, nucleus lobulation has been proposed to be important for various functions including cell transmigration (Hoffmann et al., 2007; Kolaczkowska and Kubes, 2013). Based on princeps studies performed in vivo (Bainton et al., 1971), on established tissue culture cell lines (Olins and Olins, 2004; Zhu et al., 2017), and on basic histology (Junqueira and Carneiro, 2005), the deformation of the nucleus is considered to be a late differentiation process, appearing around the metamyelocyte stage, i.e., in late precursors of neutrophils. Our study establishes instead that large nuclear invaginations are hallmarks of an early stage in the differentiation of HSCs into myeloid progenitors, and that the deformation of the nucleus in myeloid progenitors can affect the expression of genes involved in that differentiation process. This echoes similar observation in mesenchymal stem cells in which nucleus deformation has been related to the regulation of the differentiation process (Uhler and Shivashankar, 2017). Interestingly, some less striking but clear nuclear deformations consisting of narrow rifts were observed in lymphoid progenitors (Supplemental fig 1D). Their different shape implied a distinct mechanism for controlling the morphology. Nevertheless, in both types of deformations, microtubules were buried deep in the invaginations.

The role of microtubules in HSC nucleus deformation contrasts with previous observations in mesenchymal stem cells in which nucleus deformation was specifically controlled by the actin network (Uhler and Shivashankar, 2017). In HSCs, the actin network remained cortical and was not observed within the invaginations. Moreover, relaxation of the actin cortex had no impact on the deformation of nucleus in myeloid progenitors, whereas microtubule disassembly or disengagement prevented nucleus deformation. Microtubules have already been shown to apply forces on the nucleus in other systems (Lele et al., 2018), where they regulate nucleus shape (Hampoelz et al., 2011; Schreiner et al., 2015; Tariq et al., 2017), position (Cadot et al., 2012; Szikora et al., 2013), heterochromatin localization (Ramdas and Shivashankar, 2015) and envelope breakdown via dyneins (Beaudouin et al., 2002; Bolhy et al., 2011). Here also, we found that dyneins were involved in the deformation process, but the exact mechanism allowing microtubules to form bundles and produce mechanical forces on the nuclear membrane still requires further investigations.

A significant increase of nucleus volume was also associated with the process of nucleus deformation, in addition to microtubule-based forces. This expansion of the nucleus is unlikely to be driven by a softening of the nuclear membrane because the lamin content suggests it would become stiffer (Swift et al., 2013). Nucleus expansion seemed also to be independent of microtubule-based forces, because it was not prevented by microtubule disassembly (Figure 3f). The mechanism driving nucleus expansion in myeloid progenitors could depend on the regulation of nuclear osmolarity by specific ion channels (Finan et al., 2009) or be driven by chromatin decondensation due to histone modifications (Mazumder et al., 2008; Rao et al., 2007; Therizols et al., 2014). Peripheral invaginations would thus result from frustrated expansion against bundles of microtubules encircling the nucleus.

In addition to obstructing nucleus expansion, microtubules could be more directly involved in the regulation of chromatin rearrangements across the nuclear membrane. Mechanical forces can regulate chromatin organization (Miroshnikova et al., 2017), and microtubules can act specifically on chromatin across the nuclear membrane (Hampoelz et al., 2011; King et al., 2008), facilitating the pairing of homologous chromosomes (Christophorou et al., 2015), driving the positioning of chromosome territories and clustering of telomeres or centromeres beneath the centrosome (Elkouby et al., 2016; King et al., 2008). Thus, microtubules could direct chromatin reorganization during nucleus expansion in myeloid progenitors. This is supported by our observation that heterochromatin content is reduced in invaginated regions of the nucleus where microtubules accumulate. This raises the hypothesis that microtubules can specifically prevent heterochromatin anchorage or recruit euchromatin. Another hypothesis is that the regulation of heterochromatin anchoring to the nuclear periphery is independent of microtubules and sensitive to local curvature only. A weaker binding to invaginated regions would force the heterochromatin to flow away as the nucleus expands. Curvature also modulates the nuclear shuttling of transcription factors involved in stem cell fate regulation (Elosegui-Artola et al., 2017). Hence, these hypotheses could explain how perturbation of microtubules could also directly and indirectly lead to the changes in the expression of genes involved in myeloid differentiation. Therefore, for the future, it will important to investigate the localization of those genes the expression of which has been modulated by nucleus deformation, and how that relates to the shape nucleus and heterochromatin distribution, in order to further understand the mechano-regulation of HCS differentiation.

## ACKNOWLEDGEMENTS

We thank Matthieu Piel, David Pellman, Gérard Socié and Kevin Chalut for interesting discussions and critical reading of the manuscript. This work was funded by grants from the Agence Nationale pour la Recherche (ANR-14-CE11-0012, ANR-10-IHUB-0002), from the European Research Council (ERC CoG 771599), from the Emergence program of the Ville de Paris, from the “Coups d’Elan” prize of the Bettencourt-Schueller foundation, and the Schlumberger foundation for education and research. SB received PhD fellowships from the IRTELIS program of the CEA and from the Fondation pour la Recherche Medicale (grant FDT20170437071). We thank the Technological Core Facility (Plateforme Technologique de l’IRSL) of the Institut de Recherche Saint Louis, Université de Paris for technical support. The facility is supported by the Conseil Régional d’Ile-de-France, Canceropôle Ile-de-France, Université de Paris, Association Saint-Louis, Association Jean-Bernard, Fondation pour la Recherche Médicale, French National Institute for Cancer Research (InCa) and Ministère de la Recherche.

## AUTHOR CONTRIBUTIONS

S. Biedzinski performed all experiments with the help of L. Faivre, B. Vianay and S. Brunet. M. Delord analyzed the transcriptomics data. L. Blanchoin, J. Larghero, M. Théry and S. Brunet supervised the project. J. Larghero and M. Théry obtained funding for the project. M. Théry and S. Brunet conceived and directed the project, and wrote the manuscript.

## MATERIALS AND METHODS

### Cell culture

#### Blood cell harvesting and culture

Human umbilical cord blood units from normal full-term deliveries were obtained from Saint-Louis Hospital Cord Blood Bank (Paris, France) after mothers’ written informed consent, in accordance with health authorities (French Cord Blood Network, Paris, France). Mononuclear cells were collected using Ficoll separation medium (Eurobio, Courtaboeuf, France). CD34+ cells were further selected using Miltenyi Magnetically Activated Cell Sorting (MACS) columns (Miltenyi Biotech, Paris, France) according to the manufacturer’s instructions. CD34+ cells were then either put in culture or frozen at −80°C in IMDM medium (Gibco) supplemented with 10% fetal bovine serum (FBS) and 10% DMSO (WAK Chemie Medical GmbH).

#### Flow cytometry, stem and progenitor cells sorting

Freshly isolated CD34+ or thawed cells were allowed to recover overnight at 37°C in IMDM medium supplemented with10% FBS and antibiotics (Antibiotic-antimycotic, Sigma Aldrich). Cells were then labelled with fluorescent antibodies: CD45-AF700 (Clone HI30 BioLegend), CD38-PerCp5.5 (Clone HB-7 BioLegend), CD34-APC (Clone 581 BD Bioscience), CD33-PE (Clone WM-53 BD Bioscience), CD19-FITC (Clone HIB19 BD Bioscience). Sorting was performed using FACS Aria II and DIVA software (BD Bioscience). Sub-populations of stem (CD34+/CD38-), myeloid (CD38+/CD33+) and lymphoid (CD38+/CD19+) progenitors, or whole progenitors (CD34+/CD38+) for Supplementary Figure 1), were gated as shown in Figure 1B. Isolated cells were centrifuged and re-suspended in the appropriate culture medium.

#### Cell culture and drug treatments

Cells were plated at a density of 40 000 cells/cm^2^ in 96-wells plates. Cells were cultured at 37°C in a 5% CO_2_ humidified atmosphere using IMDM culture medium supplemented with antibiotics (Anti-anti, Sigma Aldrich), 10% FBS, human and SCF 100 ng/ml (Peprotech) G-CSF 10 ng/ml (Peprotech) and IL-3 20 ng/ml (Peprotech), adapted from (Donaldson et al., 2001; Faivre et al., 2016). For Figures 3, 4 and 5, in addition to control non-treated cells, cells were treated either with taxol (100 nM), blebbistatin (50 µM), nocodazole and blebbistatin (2 and 50 µM respectively), or ciliobrevin (100 µM).

#### Immunofluorescence

Cells of interest were allowed to sediment on poly-L-lysine (Sigma Aldrich) coated cover slips for 15 minutes.

For cytoskeleton staining, cells were fixed for 10 minutes with 0,5% glutaraldehyde (Sigma Aldrich) and 0,1% Triton-X100 (Sigma Aldrich) in Cytoskeleton Buffer (10 mM MES pH 6.1, 138 mM KCl, 3 mM MgCl_2_, 2 mM EGTA, 10% sucrose). Cells were permeabilized for 10 minutes in 0,1% Triton-X100 in PBS and neutralized with NaBH_4_ in PBS. Microtubules, centrosomes and microfilaments were labelled using YL1/2 rat antibody (Serotec) (1:500), rabbit anti-pericentrin (abcam) (1:2000) and CY3-phalloidin (1:100) (Sigma Aldrich), respectively.

For nucleus and chromatin staining, cells were fixed for 20 minutes with 3% paraformaldehyde (Electron Microscopy Sciences) in PBS solution, permeabilized with 0.1% Triton-X100 and 1% BSA in PBS for 20minutes and finally neutralized with NH_4_Cl in PBS. Rabbit anti-Sun2 (Sigma Aldrich; 1:1000), rabbit anti-H3K9me3 (Abcam; 1:500), monoclonal anti-H3K27me3 (Diagenode; 1:800), goat anti-lamin B (C-20; Santa Cruz; 1:800) and monoclonal anti-laminA/C (E-1; Santa Cruz; 1:500) were used.

In all cases, chromatin was labelled using DAPI (Sigma Aldrich; 5ng/ml) and samples were mounted using Mowiol solution (Sigma Aldrich).

### Data acquisition and analysis

#### Confocal Microscopy and 3D measurements

For quantifications, images were acquired using a LSM800 airy-scan or a LSM 780 confocal microscope and ZEN software (Zeiss). The objective used was a 63x APO oil immersive: an 8x digital zoom was added. Each wavelength was acquired separately with a 350nm z-step width to achieve appropriate reconstruction resolution. For nucleus shape, Image J plugin *3D viewer* was used to generate and export an isosurface of the DAPI threshold signal. The surface was then analyzed using MATLAB (Mathworks). It was imported as a mesh and smoothened using the *read_vertices_and_faces_from_obj_file script* (http://www.alecjacobson.com/weblog/?p=917, Alec Jacobson) and the *N_SmoothMesh* script (Export Voxel Data, Cyprian Lewandowski, MATLAB file exchange). The main curvatures of the surfaces were calculated using the *patchcurvature* script (Dirk-Jan Kroon, MATLAB file exchange) and corresponding areas then measured. For centrosomes, positions were detected manually in the three dimensions using pericentrin staining as a marker. An isosurface was generated and added to the nuclear reconstruction for further calculations using the *TriangleRayIntersection* script (Jaroslaw Tuszynski, MATLAB file exchange). Images were generated with MATLAB (Mathworks). Data were plotted using Prism software (GraphPad Software, Inc).

#### Chromatin image analysis

Images were acquired using a LSM 780 confocal microscope and ZEN software (Zeiss). The objective used was a 63x oil immersive (model): an 8x digital zoom was added. Single planar slices were manually selected, post-acquisition, to maximize deformation in a 2D plane. The contour of the H3K9me3 signal was manually drawn on Image J in order to obtain precise line scans. Curvature values were obtained based on an automated shape detection, to improve spatial resolution compared with manual detection. Linescan values and polygon vertices were extracted from Image J and analyzed with MATLAB. Straight views of the line selection were obtained with Image J *straighten* function. All data are plotted with Prism software (GraphPad Software, Inc).

#### Transcriptomic analysis

RNA quantification and quality control was performed using the HT RNA Pico sensitivity LabChip Kit and the Caliper LabChip Microfluidics System (Perkin Elmer). For each sample, 3 ng of total RNA was amplified, labeled, and fragmented using GeneChip WT Pico (ThermoFisher Scientific). Each sample was hybridized onto Human Clariom D (ThermoFisher Scientific), washed, and stained with the Affymetrix® Fluidics Station 450. Array scanning was performed with the Affymetrix® GeneChip Scanner 3000 7G using the Command Console software (ThermoFisher Scientific) and then analyzed using the Affymetrix® rma-sketch routine. Differential expression between condition were analysed using the limma routine implemented in the limma R library (Smyth, 2004).

#### Statistical analysis

All statistical analyses i.e Mann-Whitney tests were performed with Prism software (GraphPad Software, inc).

---

**Supp. Figure 1.**
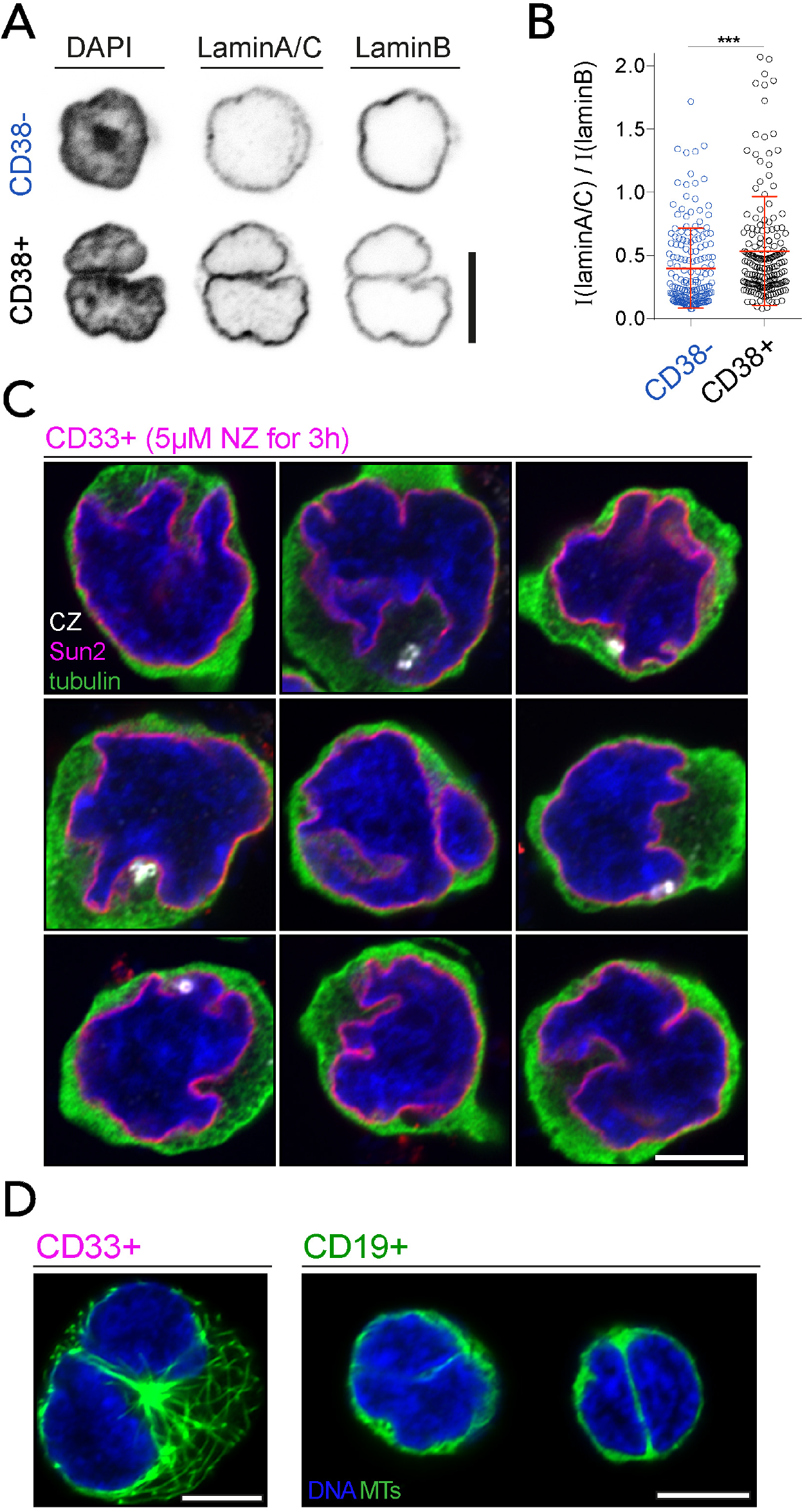
CD33+ nucleus undergoes plastic deformations. A) Lamin composition of stem *vs* progenitor cells. Chromatin, lamin A/C and lamin B immunostaining in representative HSCs (CD38-, blue) versus progenitors (CD38+, black). Inverted images of the equatorial Z stack are presented. Scale bar: 5 µm. B) The lamin (A/C) / lamin B ratio is significantly higher in CD38+ (n=191) than CD38-cells (n=169; ***: p<0.001 Mann Whitney test), indicative of a nuclear envelope stiffening upon differentiation. C) Gallery of selected Z stacks of representative CD33+ cells treated for 3 hours with 5 µM nocodazole. Nucleus envelope (Sun2 immuno-staining) is shown in magenta, chromatin in blue, and centrosome in white. The tubulin signal in green has been enhanced to show the lack of polymerized microtubules. Scale bar 5 µm. D) Microtubules are associated with CD33+ nuclear invaginations and CD19+ nuclear rifts. Z stacks of representative cells are shown. Microtubules are shown in green and chromatin in blue. Scale bar 5 µm.

**Supp. Figure 2.**
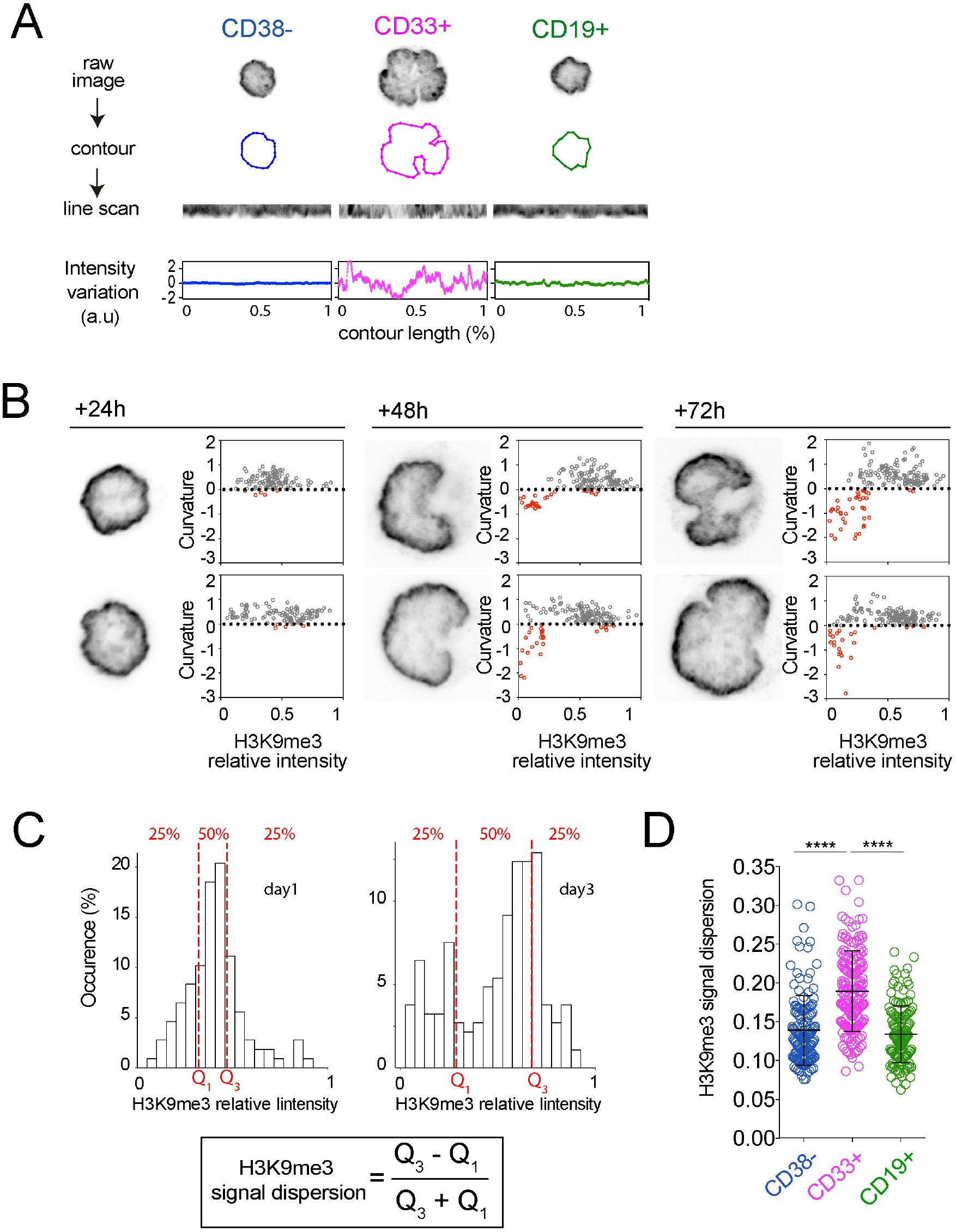
H3K9me3 distribution in CD38-, CD33+ and CD19+ isolated cells. A) For each cell, the equatorial Z stack of the raw DAPI image is used to extract the nucleus contour (middle row) and measure H3K9Me3 intensity (in arbitrary units [a.u]) along the nucleus contour (lower row). B) For each time point, inverted images of equatorial Z stacks of H3K9me3 (left panel) of two representative nuclei are presented. For each cell, local curvature of the nucleus envelope is plotted as a function of H3K9Me3 local intensity at the indicated time points (right panel). Positive and negative and curvatures are shown in grey and red, respectively. Null curvature is highlighted as a dashed line. C) Histograms show the values of H3K9me3 signal intensity along the nucleus contour of two cell representatives of high and low signal dispersion, respectively. The distribution of values can be segmented in quartiles. H3K9me3 signal-intensity dispersion is measured as the quartile coefficient of dispersion (Q disp = (Q3-Q1)/(Q3+Q1), where Q1 and Q3 are the first and the third quartiles, respectively, of the H3K9me3 intensity distribution on the nucleus contour. B) The dispersion parameter is higher in CD33+ (n=124, 3 donors) than in CD38-(n=137, 3donors) and CD19+ (n=132/ 3 donors) cells (****; p<0.0001. Mann Whitney test) indicating that H3K9me3 is homogenously distributed at the nucleus periphery in CD38- and CD19+ cells, but is heterogeneous at the nucleus periphery in CD33+ cells.

**Supplementary Table 1.**
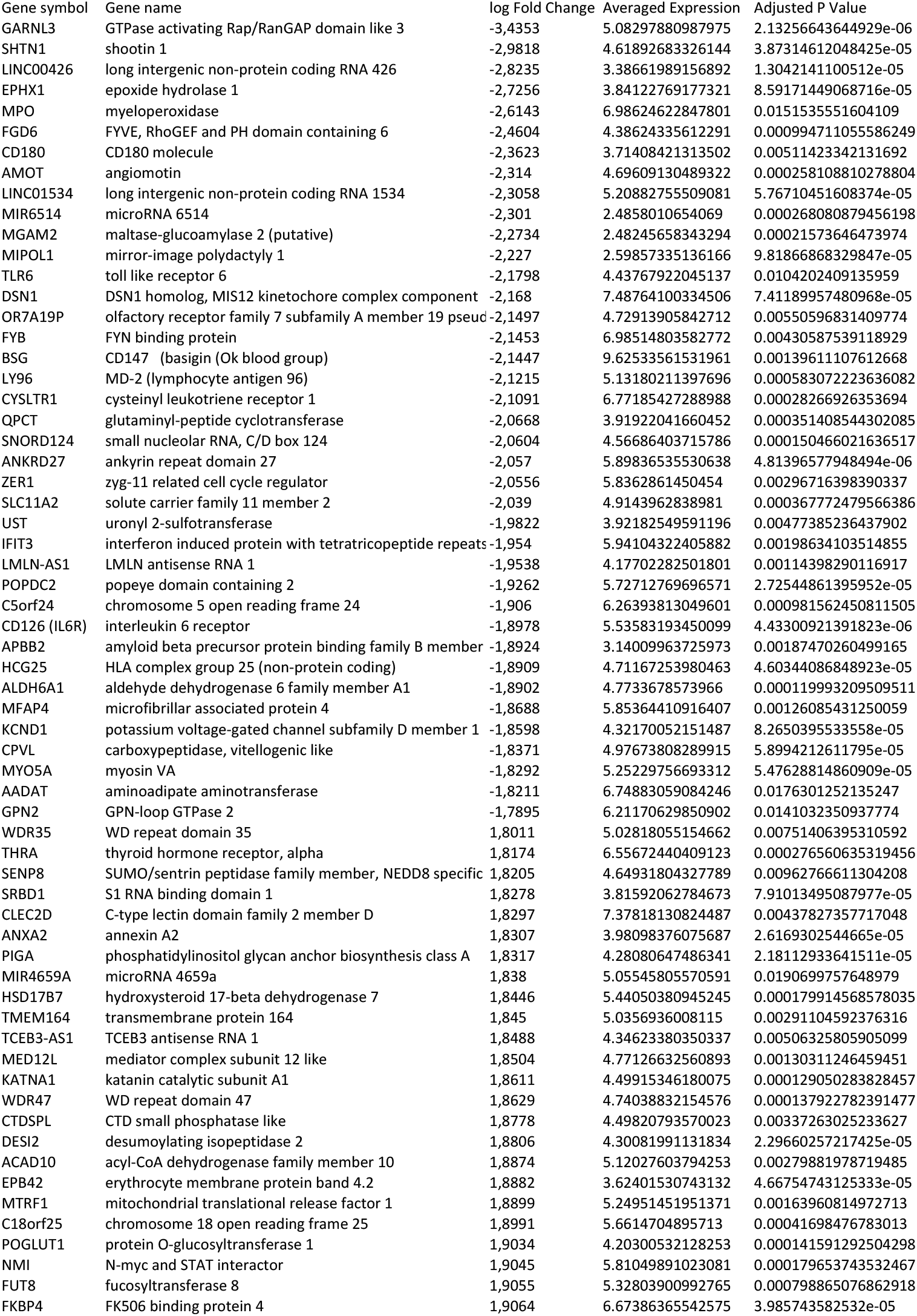
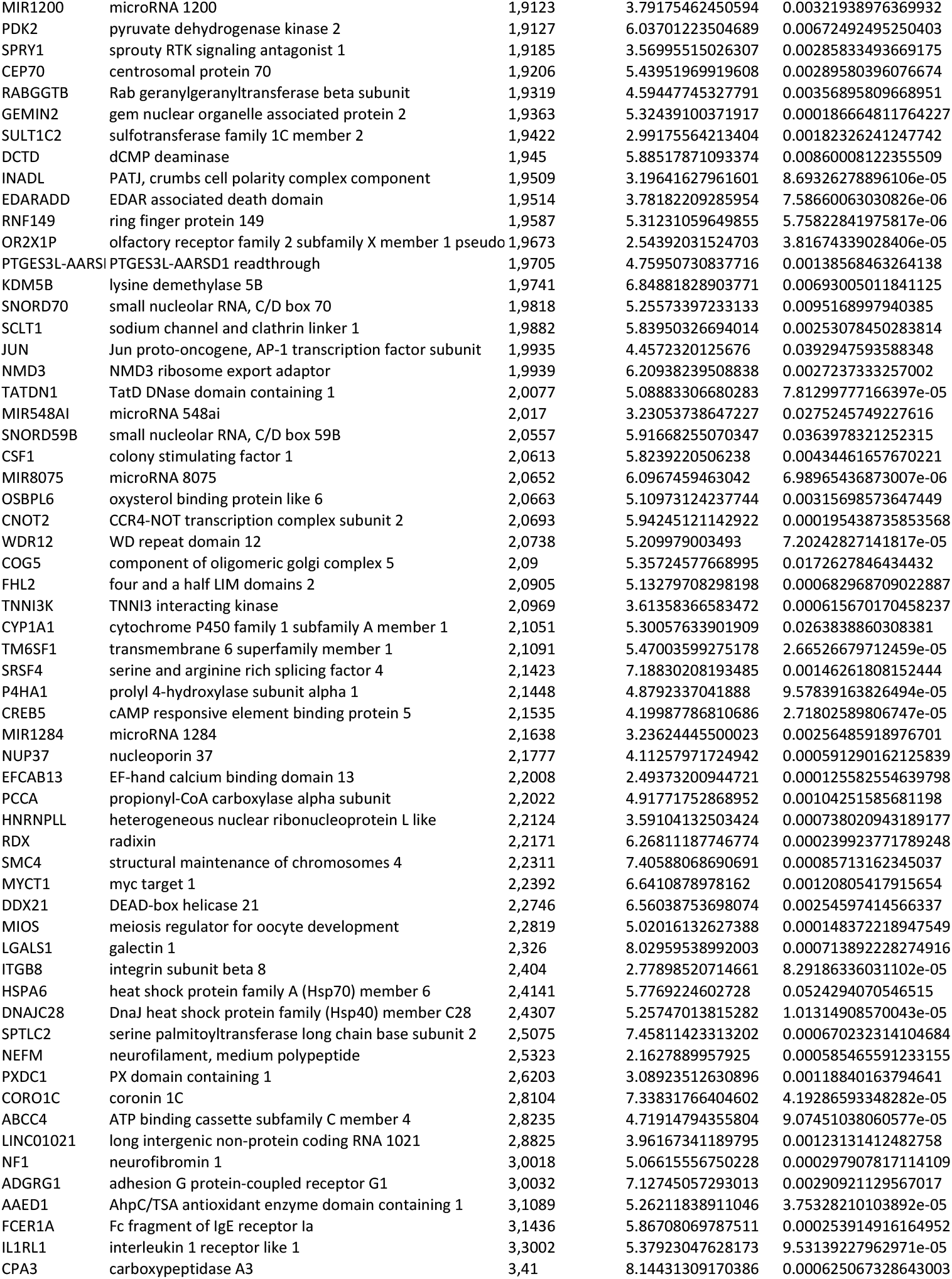
List of the significantly up- and down-regulated genes in ciliobrevin D treated cells with respect to non-treated cells. Gene symbols, ID full names and log2 fold changes values are indicated.

## REFERENCES

Alam, S.G., Zhang, Q., Prasad, N., Li, Y., Chamala, S., Kuchibhotla, R., Kc, B., Aggarwal, V., Shrestha, S., Jones, A.L., et al. (2016). The mammalian LINC complex regulates genome transcriptional responses to substrate rigidity. Nat. Publ. Gr. 1–11.

Bainton, D.F., Ullyot, J.L., and Farquhar, M.G. (1971). The development of neutrophilic polymorphonuclear leukocytes in human bone marrow: origin and content of azurophil and specific granules. J. Exp. Med. 134, 907–934.

Bannister, A.J., Zegerman, P., Partridge, J.F., Miska, E.A., Thomas, J.O., Allshire, R.C., and Kouzarides, T. (2001). Selective recognition of methylated lysine 9 on histone H3 by the HP1 chromo domain. Nature 410, 120–124.

Beaudouin, J., Gerlich, D., Daigle, N., Eils, R., and Ellenberg, J. (2002). Nuclear Envelope Breakdown Proceeds by Microtubule-Induced Tearing of the Lamina. Cell 108, 83–96.

Becker, J.S., Nicetto, D., and Zaret, K.S. (2016). H3K9me3-Dependent Heterochromatin: Barrier to Cell Fate Changes. Trends Genet. 32, 29–41.

Bolhy, S., Bouhlel, I., Dultz, E., Nayak, T., Zuccolo, M., Gatti, X., Vallee, R., Ellenberg, J., and Doye, V. (2011). A Nup133-dependent NPC-anchored network tethers centrosomes to the nuclear envelope in prophase. J. Cell Biol. 192, 855–871.

Buxboim, A., Swift, J., Irianto, J., Spinler, K.R., Dingal, P.C.D.P., Athirasala, A., Kao, Y.-R.C., Cho, S., Harada, T., Shin, J., et al. (2014). Matrix Elasticity Regulates Lamin-A,C Phosphorylation and Turnover with Feedback to Actomyosin. Curr. Biol. 24, 1909–1917.

Cadot, B., Gache, V., Vasyutina, E., Falcone, S., Birchmeier, C., and Gomes, E.R. (2012). Nuclear movement during myotube formation is microtubule and dynein dependent and is regulated by Cdc42, Par6 and Par3. EMBO Rep. 13, 741–749.

Carvalho, L.O., Aquino, E.N., Neves, A.C.D., and Fontes, W. (2015). The Neutrophil Nucleus and Its Role in Neutrophilic Function. J. Cell. Biochem. 116, 1831–1836.

Christophorou, N., Rubin, T., Bonnet, I., Piolot, T., Arnaud, M., and Huynh, J.-R. (2015). Microtubule-driven nuclear rotations promote meiotic chromosome dynamics. Nat. Cell Biol.

Crane, G.M., Jeffery, E., and Morrison, S.J. (2017). Adult haematopoietic stem cell niches. Nat. Publ. Gr. 17, 573–590.

Djeghloul, D., Kuranda, K., Kuzniak, I., Barbieri, D., Naguibneva, I., Choisy, C., Bories, J.-C., Dosquet, C., Pla, M., Vanneaux, V., et al. (2016). Age-Associated Decrease of the Histone Methyltransferase SUV39H1 in HSC Perturbs Heterochromatin and B Lymphoid Differentiation. Stem Cell Reports 6, 970–984.

Donaldson, C., Bradley, B., and Hows, J. (2001). The CD34+CD38neg population is significantly increased in haemopoietic cell expansion cultures in serum-free compared to serum-replete conditions: Dissociation of phenotype and function. Bone Marrow Transplant. 27, 365–371.

Dupont, S., Morsut, L., Aragona, M., Enzo, E., Giulitti, S., Cordenonsi, M., Zanconato, F., Digabel, J. Le, Forcato, M., Bicciato, S., et al. (2011). Role of YAP / TAZ in mechanotransduction.

Ehrlich, P., and Lazarus, A. (1900). Histology of Blood, normal and pathological (London: Cambridge University Press).

Elkouby, Y.M., Jamieson-Lucy, A., and Mullins, M.C. (2016). Oocyte Polarization Is Coupled to the Chromosomal Bouquet, a Conserved Polarized Nuclear Configuration in Meiosis. PLOS Biol. 14, e1002335.

Elosegui-Artola, A., Andreu, I., Beedle, A.E.M., Lezamiz, A., Uroz, M., Kosmalska, A.J., Oria, R., Kechagia, J.Z., Rico-Lastres, P., Le Roux, A.L., et al. (2017). Force Triggers YAP Nuclear Entry by Regulating Transport across Nuclear Pores. Cell 171, 1397–1410.e14.

Faivre, L., Parietti, V., Siñeriz, F., Chantepie, S., Gilbert-sirieix, M., Albanese, P., Larghero, J., and Vanneaux, V. (2016). In vitro and in vivo evaluation of cord blood hematopoietic stem and progenitor cells amplified with glycosaminoglycan mimetic. Stem Cell Res. Ther. 1–10.

Finan, J.D., Chalut, K.J., Wax, A., and Guilak, F. (2009). Nonlinear Osmotic Properties of the Cell Nucleus. Ann. Biomed. Eng. 37, 477–491.

Firestone, A.J., Weinger, J.S., Maldonado, M., Barlan, K., Langston, L.D., O’Donnell, M., Gelfand, V.I., Kapoor, T.M., and Chen, J.K. (2012). Small-molecule inhibitors of the AAA+ ATPase motor cytoplasmic dynein. Nature 484, 125–129.

Gupta, S., Marcel, N., Sarin, A., and Shivashankar, G. V. (2012). Role of Actin Dependent Nuclear Deformation in Regulating Early Gene Expression. PLoS One 7, e53031.

Hampoelz, B., Azou-Gros, Y., Fabre, R., Markova, O., Puech, P.-H., and Lecuit, T. (2011). Microtubule-induced nuclear envelope fluctuations control chromatin dynamics in Drosophila embryos. Development 3386, 3377–3386.

Hoffmann, K., Sperling, K., Olins, A.L., and Olins, D.E. (2007). The granulocyte nucleus and lamin B receptor: Avoiding the ovoid. Chromosoma 116, 227–235.

Junqueira, L.C., and Carneiro, J. (2005). Basic histology, text and atlas (McGraw-Hill Medical).

Khatau, S.B., Hale, C.M., Stewart-hutchinson, P.J., Patel, M.S., Stewart, C.L., Searson, P.C., Hodzic, D., and Wirtz, D. (2009). A perinuclear actin cap regulates nuclear shape. PNAS.

Kilian, K. a, Bugarija, B., Lahn, B.T., and Mrksich, M. (2010). Geometric cues for directing the differentiation of mesenchymal stem cells. Proc. Natl. Acad. Sci. U. S. A. 107, 4872–4877.

King, M.C., Drivas, T.G., and Blobel, G. (2008). A Network of Nuclear Envelope Membrane Proteins Linking Centromeres to Microtubules. Cell 134, 427–438.

Kolaczkowska, E., and Kubes, P. (2013). Neutrophil recruitment and function. Nat. Rev. Immunol. 13, 159–175.

Kolodney, M.S., and Elson, E.L. (1995). Contraction due to microtubule disruption is associated with increased phosphorylation of myosin regulatory light chain. PNAS 92, 10252–10256.

Lele, T.P., Dickinson, R.B., and Gundersen, G.G. (2018). Mechanical principles of nuclear shaping and positioning. J. Cell Biol. 217, 3330–3342.

Makhija, E., Jokhun, D.S., and Shivashankar, G. V. (2015). Nuclear deformability and telomere dynamics are regulated by cell geometric constraints. Proc. Natl. Acad. Sci. 201513189.

Mattout, A., Cabianca, D.S., and Gasser, S.M. (2015). Chromatin states and nuclear organization in development — a view from the nuclear lamina. Genome Biol. 16, 174.

Mazumder, A., Roopa, T., Basu, A., Mahadevan, L., and Shivashankar, G. V (2008). Dynamics of Chromatin Decondensation Reveals the Structural Integrity of a Mechanically Prestressed Nucleus. Biophys. J. 95, 3028–3035.

McBeath, R., Pirone, D.M., Nelson, C.M., Bhadriraju, K., and Chen, C.S. (2004). Cell shape, cytoskeletal tension, and RhoA regulate stem cell lineage commitment. Dev. Cell 6, 483–495.

Miralles, F., Posern, G., Zaromytidou, A., and Treisman, R. (2003). Actin Dynamics Control SRF Activity by Regulation of Its Coactivator MAL. Cell 113, 329–342.

Miroshnikova, Y.A., Nava, M.M., and Wickström, S.A. (2017). Emerging roles of mechanical forces in chromatin regulation. J. Cell Sci. 130, 2243–2250.

Närvä, E., Stubb, A., Guzmán, C., Blomqvist, M., Balboa, D., Lerche, M., Saari, M., Otonkoski, T., and Ivaska, J. (2017). A Strong Contractile Actin Fence and Large Adhesions Direct Human Pluripotent Colony Morphology and Adhesion. Stem Cell Reports 9, 67–76.

Notta, F., Zandi, S., Takayama, N., Dobson, S., Gan, O.I., Wilson, G., Kaufmann, K.B., McLeod, J., Laurenti, E., Dunant, C.F., et al. (2016). Distinct routes of lineage development reshape the human blood hierarchy across ontogeny. Science (80-.). 351.

Olins, A.L., and Olins, D.E. (2004). Cytoskeletal influences on nuclear shape in granulocytic HL-60 cells. BMC Cell Biol. 5, 1–18.

Orkin, S.H., and Zon, L.I. (2008). Hematopoiesis: An Evolving Paradigm for Stem Cell Biology. Cell 132, 631–644.

Paluch, E., Piel, M., Prost, J., Bornens, M., and Sykes, C. (2005). Cortical actomyosin breakage triggers shape oscillations in cells and cell fragments. Biophys. J. 89, 724–733.

Pinho, S., and Frenette, P.S. (2019). Haematopoietic stem cell activity and interactions with the niche. Nat. Rev. Mol. Cell Biol.

Ramdas, N.M., and Shivashankar, G. V. (2015). Cytoskeletal Control of Nuclear Morphology and Chromatin Organization. J. Mol. Biol. 427, 695–706.

Rao, J., Bhattacharya, D., Banerjee, B., Sarin, A., and Shivashankar, G. V (2007). Trichostatin-A induces differential changes in histone protein dynamics and expression in HeLa cells. 363, 263–268.

Salina, D., Bodoor, K., Eckley, D.M., Schroer, T.A., Rattner, J.B., Burke, B., Tn, A., and Hall, M. (2002). Cytoplasmic Dynein as a Facilitator of Nuclear Envelope Breakdown. 108, 97–107.

Schreiner, S.M., Koo, P.K., Zhao, Y., Mochrie, S.G.J., and King, M.C. (2015). The tethering of chromatin to the nuclear envelope supports nuclear mechanics. Nat. Commun. 6, 7159.

Shin, J.-W., Swift, J., Ivanovska, I., Spinler, K.R., Buxboim, A., and Discher, D.E. (2013). Mechanobiology of bone marrow stem cells: from myosin-II forces to compliance of matrix and nucleus in cell forms and fates. Differentiation 86, 77–86.

Shin, J., Buxboim, A., Spinler, K.R., Swift, J., Christian, D.A., Hunter, C.A., Léon, C., Gachet, C., Dingal, P.C.D.P., Ivanovska, I.L., et al. (2014). Contractile forces sustain and polarize hematopoiesis from stem and progenitor cells. Cell Stem Cell 14, 81–93.

Smyth, G.K. (2004). Linear Models and Empirical Bayes Methods for Assessing Differential Expression in Microarray Experiments. Stat. Appl. Genet. Mol. Biol. 3, 1–25.

Swift, J., Ivanovska, I.L., Buxboim, A., Harada, T., Dingal, P.C.D.P., Pinter, J., Pajerowski, J.D., Spinler, K.R., Shin, J.-W., Tewari, M., et al. (2013). Nuclear lamin-A scales with tissue stiffness and enhances matrix-directed differentiation. Science 341, 1240104.

Szikora, S., Gaspar, I., and Szabad, J. (2013). ‘Poking’ microtubules bring about nuclear wriggling to position nuclei. J. Cell Sci. 126, 254–262.

Tajik, A., Zhang, Y., Wei, F., Sun, J., Jia, Q., Zhou, W., Singh, R., Khanna, N., Belmont, A.S., and Wang, N. (2016). Transcription upregulation via force-induced direct stretching of chromatin. Nat. Mater. 15, 1287–1296.

Tariq, Z., Zhang, H., Chia-Liu, A., Shen, Y., Gete, Y., Xiong, Z.-M., Tocheny, C., Campanello, L., Wu, D., Losert, W., et al. (2017). Lamin A and microtubules collaborate to maintain nuclear morphology. Nucleus 8, 1–14.

Therizols, P., Illingworth, R.S., Courilleau, C., Boyle, S., Wood, A.J., and Bickmore, W. a (2014). Chromatin decondensation is sufficient to alter nuclear organization in embryonic stem cells. Science 346, 1238–1242.

Ugarte, F., Sousae, R., Cinquin, B., Martin, E.W., Krietsch, J., Sanchez, G., Inman, M., Tsang, H., Warr, M., Passegu??, E., et al. (2015). Progressive chromatin condensation and H3K9 methylation regulate the differentiation of embryonic and hematopoietic stem cells. Stem Cell Reports 5, 728–740.

Uhler, C., and Shivashankar, G. V. (2017). Regulation of genome organization and gene expression by nuclear mechanotransduction. Nat. Rev. Mol. Cell Biol.

Velten, L., Haas, S.F., Raffel, S., Blaszkiewicz, S., Islam, S., Hennig, B.P., Hirche, C., Lutz, C., Buss, E.C., Nowak, D., et al. (2017). Human haematopoietic stem cell lineage commitment is a continuous process. Nat. Cell Biol. 19, 271–281.

Versaevel, M., Grevesse, T., and Gabriele, S. (2012). Spatial coordination between cell and nuclear shape within micropatterned endothelial cells. Nat. Commun. 3, 671.

Watkins, N.A., Foad, N.S., Garner, S.F., Jolley, J., Koch, K., Macaulay, I.C., Morley, S.L., Rendon, A., Taylor, N., Winzer, T., et al. (2009). A HaemAtlas: Characterizing gene expression in differentiated human blood cells. Blood 113, 1–10.

Zhu, Y., Gong, K., Denholtz, M., Chandra, V., Kamps, M.P., Alber, F., and Murre, C. (2017). Comprehensive characterization of neutrophil genome topology. Genes Dev. 31, 141–153.

